# Species- and Topic-aware Representation Learning for Antimicrobial Peptide Discovery

**DOI:** 10.64898/2026.05.28.728246

**Authors:** Sarala Padi, Kinjal Mondal, Navleen Kaur, David Hoogerheide, Frank Heinrich, Mihaela Mihailescu, Jeffery B. Klauda, Antonio Cardone, Walid Keyrouz

## Abstract

Antimicrobial resistance poses a major global health challenge, necessitating efficient strategies to discover potent antimicrobial peptides (AMPs). While recent generative models can produce many candidate sequences, experimentally validating all generated peptides in wet labs is impractical due to the high costs and time involved in such measurements. As a result, there is a strong demand for accurate predictions of peptide efficacy, typically measured as the minimum inhibitory concentration (MIC). We introduce STAMP, a framework for Species- and Topic-aware Representation Learning in AMP Discovery. This unified machine learning framework allows for cross-species predictions of AMP activity. STAMP integrates protein language model embeddings with species conditioning and topic-aware representations that capture sequence-level patterns, enabling generalizable predictions across multiple bacterial species within a single model. We evaluated STAMP on three benchmark datasets, which include two previously published datasets and a newly curated dataset derived from DBAASP, addressing duplicates and inconsistencies systematically. STAMP achieved strong predictive performance across these datasets, demonstrating a Pearson correlation coefficient (PCC) of 0.837 and an R^2^ of 0.70, outperforming several baseline models. Importantly, we further validated our prediction model using peptides that were experimentally tested for their antimicrobial activity against *E*.*coli*. and *S*.*epidermidis* bacteria, demonstrating its real-world applicability. Furthermore, residue-level importance analyses provide insights into the sequence determinants governing antimicrobial activity. Together, these results establish STAMP as a scalable framework for MIC prediction and an effective computational tool for accelerating AMP discovery and optimization.

## Introduction

Antibiotics play a critical role in treating life-threatening infections by targeting and eliminating specific bacteria [1]. However, their overuse can lead to side effects and contribute to antimicrobial resistance (AR), where bacteria and fungi evolve to resist the drugs meant to kill them [2]. The COVID-19 pandemic highlighted the dangers of inappropriate antibiotic use and extended hospital stays [3, 4, 5]. Overuse and misuse of antibiotics in humans, animals, and agriculture, along with gaps in infection prevention and drug development, are driving the AR crisis [6]. As traditional antibiotic development lags and resistance spreads, there is an urgent need for alternative strategies. Antimicrobial peptides (AMPs) are an alternative to antibiotics and form a diverse class of naturally occurring molecules that directly kill bacteria, yeasts, fungi, viruses, and even cancer cells [7]. AMPs show promise in disrupting bacterial membranes and use multi-target mechanisms, making it difficult for AR to develop. This approach could provide a new class of therapeutics essential for managing AR and ensuring effective treatment for infections in the future [8, 7].

Recent advances in the design of AI-driven AMPs clearly demonstrate a decisive trend toward integrating generative modeling frameworks with the prediction of MIC as a core task for screening and optimization. Popular generative models include GAN- and GPT-based frameworks [9, 10], along with feature-driven models like DLFea4AMPGen [11], diffusion models such as AMPGen [12], and variational autoencoders like PepVAE [13]. Furthermore, innovative hybrid pipelines that merge generative models with molecular dynamics simulations and wet-lab testing play a critical role in the development process [14, 15]. Collectively, these findings illustrate a compelling consensus: regardless of the specific architectural approaches—whether they are GANs, VAEs, diffusion models, or large language models—modern AMP discovery pipelines fundamentally rely on determining the minimum inhibitory concentration (MIC) as a quantitative link connecting sequence generation to experimental validation.

The MIC is a key measure for evaluating antibiotic resistance, measuring the lowest concentration of an antibiotic that inhibits bacterial growth using agar or broth methods [16, 17, 18, 19]. While traditional methods like broth micro-dilution and disk diffusion assays are considered the gold standard, they are labor-intensive, time-consuming, and reliant on the growth of the bacteria, often requiring 18 to 24 hours for results. This can delay critical treatment decisions [16, 17, 18]. With the vast number of sequences generated by AI models, it would be impractical to measure all of them for validation. Therefore, effective filtering and selection of the most promising AMPs are essential for efficiently directing resources toward the best candidates [17, 20, 21]. Therefore, building a predictive model that estimates the MIC directly from protein or genomic sequences provides a rapid and scalable alternative. This approach not only saves time and resources but also increases the likelihood of identifying effective AMPs for further investigation.

Predicting AMP activity is a significant challenge in antimicrobial discovery due to species-specific variability in efficacy. Identical peptide sequences can exhibit substantial differences in MIC across bacteria, emphasizing the need for species-aware predictive models [22]. Recent methods that combine peptide sequence embeddings with genomic descriptors show promise but rely heavily on genome availability and prior knowledge, limiting their use with new peptides and under-characterized species [23, 24, 25].

Topic model [26] is a well-established method in natural language processing (NLP) domain to extract hidden patterns in text data by transforming high-dimensional data into compact representations. Here, we introduce Species- and Topic-aware Representation Learning for AMP discovery (STAMP), sequence-centric modeling framework that conditions MIC prediction on peptide embeddings alongside compact representations of species and latent sequence patterns. By leveraging Evolutionary Scale Modeling (ESM2) [27], STAMP extracts context-aware, order-sensitive embeddings that capture biologically meaningful motifs without relying on genome-level features. Species information is incorporated through lightweight categorical encodings, while topic embeddings enable scalable multimodal learning with improved interpretability.

Building on our prior work on C-terminal poly-arginine segments [28], we further evaluate STAMP on peptides tested against *E. coli*. and *S. epidermidis*, including native, truncated, and cationic tail–modified variants. STAMP accurately captures the resulting shifts in MIC, demonstrating its ability to learn physicochemical determinants of pathogen-specific antimicrobial activity.

### Related Work: MIC Prediction for Antimicrobial Peptides

Recent advancements in machine learning (ML) and deep learning (DL) have significantly enhanced the prediction of AR phenotypes and MICs from both genomic and peptide sequences. Genome-based models trained on extensive datasets, such as thousands of Salmonella isolates, demonstrate that whole-genome features can effectively predict MICs across various antibiotics while identifying informative resistance-associated regions [21]. Concurrently, peptide-focused approaches utilizing regression models and feature engineering, including Random Forest and deep learning architectures, show moderate-to-strong correlations with experimental MIC data, indicating their potential in estimating AMP efficacy [29, 24]. Advanced ensemble deep learning frameworks that integrate sequence-derived and pathogen-specific genomic features have further improved MIC predictions, particularly against priority bacterial strains [24]. Reviews highlight that despite their promising predictive abilities, many AI-driven AR studies remain in the early translational phase, facing challenges like feature selection, data heterogeneity, model generalizability, and clinical deployment [30, 31].

Moreover, Yan et.al leveraged deep learning architectures to enhance AMP activity prediction, overcoming challenges such as data imbalance and cyclic peptides. Convolutional neural networks (CNNs) have been employed to improve short-length AMP predictions through optimal amino acid composition features, achieving high accuracy and facilitating genome-wide screening for effective AMPs [32]. Multi-branch convolutional networks with attention mechanisms have demonstrated superior MIC prediction performance in comparison to traditional machine learning models [33]. Additionally, graph neural networks have captured k-mer-level genomic similarities, providing insights into resistance determinants [34]. The adoption of protein language models (PLMs) further enhances AMP-related tasks by yielding contextualized residue embeddings through large-scale pretraining, showing improved classification and activity prediction outcomes [35, 36].

All existing MIC prediction models encounter several critical limitations. Traditional machine learning and early deep learning approaches heavily depend on handcrafted physicochemical descriptors, resulting in limited generalizability, particularly when applied to peptides outside the training distribution [37]. While sequence-based deep learning models enhance representation learning, they frequently overlook species-dependent variability in antimicrobial activity.

Moreover, although species-aware models represent a significant step forward, their reliance on predefined genomic descriptors restricts applicability. Genomic features are often linked to specific bacterial species and may not be readily obtainable for newly generated peptides or hypothetical target organisms. This limitation hampers the generalizability of these models in de novo peptide design or exploratory antimicrobial discovery scenarios where genomic context is uncertain or unavailable [23]. Additionally, many existing approaches treat MIC prediction as a conventional regression problem despite the ordinal and dilution-based nature of MIC measurements; experimental MIC values are generally reported in discrete concentration ranges and exhibit inherent assay variability not captured by standard regression losses [37]. A further concern arises from the lack of systematic analysis regarding sequence order perturbations, raising questions about whether current models effectively utilize true biological order-dependent signals rather than merely global compositional statistics [38].

These limitations motivate the development of an species-aware protein language modeling framework that enables robust and scalable MIC prediction without relying on species-specific genomic features. Our approach integrates ESM-2 embeddings with species- and topic-aware representations to capture both fine-grained residue-level context and higher-order antimicrobial activity patterns derived from dataset-level structure. The combination of contextual sequence embeddings and latent themes from topic embeddings provides complementary information that enhances predictive accuracy. By incorporating lightweight species conditioning alongside topic embeddings, the proposed framework supports scalable, species-aware MIC prediction across diverse microbial populations, facilitating antimicrobial peptide design and therapeutic optimization.

## Materials and Methods

### Datasets

In this paper, we focus on two benchmark datasets that have been published in the literature for research purposes. The first dataset was introduced by Daehun Bae et al. [23]. The authors collected a comprehensive corpus of approximately 1.7 million peptide sequences for the masked language model (MLM) pre-training of ESM-2. This pre-training enhances peptide representation learning specifically for short antimicrobial sequences rather than entire proteins. For the species-aware MIC regression task, the fine-tuning dataset was curated from DBAASP v3, resulting in 9,992 peptide–species pairs spanning 35 bacterial species. Each pair includes experimentally measured MIC values in micromolar (µM), which were log10-transformed for regression analysis. Only species with at least 60 associated peptides and sequences ranging from 5 to 50 amino acids were included. The dataset was divided by species into training, validation, and test sets using an 80:10:10 ratio across the 35 species used for evaluation (see Table 1). Bae et al. proposed the LLAMP model for MIC prediction, which was evaluated on the hold-out data. The performance of MIC prediction varied across species, with over half showing Pearson correlation coefficients above 0.7, highlighting the challenge of capturing species-specific activity with limited data. The authors also used the model to screen approximately 5.5 million candidate peptides from PeptideAtlas, targeting low predicted MIC values (less than 10 µM) across both Gram-positive and Gram-negative targets. They applied physicochemical and structural filters before selecting the top experimental candidates.

**Table 1.** Datasets, species and number of sequences used for model training, validation and testing evaluations.

| Database | Species | No: of Sequences used for Model Training and Evaluations |  |  |
| --- | --- | --- | --- | --- |
|  |  | Train | Test | Validation |
| Dataset A (Bae et. al [23]) | *E.coli.* | 6095 | 749 | 762 |
|  | *S.aureus.* | 5518 | 695 | 695 |
|  | *P.aeruginosa.* | 3978 | 509 | 507 |
|  | *S.Epidermidis.* | 1458 | 190 | 178 |
|  | **Total** | 29394 | 3741 | 3683 |
| Dataset B (Chung et. al [24]) | *E.coli.* | 2443 | 764 | 611 |
|  | *S.aureus.* | 1692 | 529 | 423 |
|  | *P.aeruginosa.* | 1572 | 492 | 394 |
|  | **Total** | 5707 | 1785 | 1428 |
| Our Curated Dataset | *E.coli.* | 5296 | 654 | 589 |
|  | *S.aureus.* | 4557 | 563 | 507 |
|  | *P.aeruginosa.* | 3752 | 464 | 417 |
|  | *S.Epidermidis.* | 1646 | 204 | 182 |
|  | **Total** | 15251 | 1885 | 1695 |

The second dataset we consider for evaluating the MIC prediction model was published by Chung et al. [24]. This study focuses on MIC prediction for AMPs targeting three priority bacterial strains: Staphylococcus aureus ATCC 25923, Escherichia coli ATCC 25922, and Pseudomonas aeruginosa ATCC 27853. The authors employed regression models that integrate both peptide and genomic sequence features. The dataset was assembled from multiple AMP repositories, including DBAASP, dbAMP, and DRAMP. After rigorous preprocessing to include only peptides of typical AMP length (6–40 amino acids) and to eliminate duplicates, the dataset resulted in 8,920 unique AMP–strain combinations distributed across training, validation, and independent testing sets. For instance, the independent test set comprised 1,785 AMP sequences, while the final training set included 5,707 examples aggregated across the three strains (refer to Table 1). The MIC values were analyzed on a logarithmic scale (log µM) to account for the wide variability in activity, and peptides were classified into low, medium, and high MIC ranges to illustrate distributional properties. The ensemble model combined predictions from bi-directional long short-term memory (BiLSTM), convolutional neural network (CNN), and multi-branch deep learning architectures, achieving strong Pearson correlations between predicted and actual MIC values (approximately 0.75–0.80 depending on the strain).

### Our curated dataset (DBAASP-v3)

Every year, AMP repositories are updated with new sequences and measurements. To obtain the most current datasets, we applied our data processing methods to curate the full Database of Antimicrobial Activity and Structure of Peptides (DBAASP) dataset. During this process, we noticed inconsistencies in reporting MIC values, units, and ranges. Some MIC values were given as a range (e.g., x-y), where x is less than y, but this assumption does not hold for all measurements. There are instances where MIC is incorrectly reported as x-y with x greater than y. Additionally, some MIC values are defined as greater than a given value x. Generally these values are converted to 2x for cases of “greater than” and x/2 for “less than.” Some papers treat x uniformly, regardless of whether it is designated as “greater than” or “less than.” The units used to report MIC values also vary [39, 40].

To address these inconsistencies, MIC values, often reported as ranges or non-numeric strings across various databases, were standardized to a single concentration in *µM* following protocols consistent with the DBAASP API. Specifically, singular values (*x*) were recorded directly, while the maximum value was selected from ranges (*x* − *y*) to represent the upper bound of inhibitory activity. For measurements reported with standard deviations (*x* ± *y*), the mean (*x*) was utilized. In instances involving inequality symbols (e.g., > *x*, ≤ *x*) or comparative ranges, all symbols were removed and the highest indicated numerical value was retained. This standardization resulted in a uniform quantitative dataset optimized for computational analysis.

The DBAASP dataset contains multiple entries for identical peptide sequences with varying MIC values due to differences in experimental conditions and measurement protocols. To ensure a well-defined mapping between sequence and activity and to avoid potential data leakage and label inconsistency, unique peptide sequences were retained for model training and evaluation. We collected sequences from all species categories, totaling 43, 477, of which we identified 18, 831 unique peptide sequences specific to four target species, as listed in Table 1.

Regarding the database division into training, testing, and validation sets, both Daehun Bae et al. [23] and Chung et al. [24] used a random 80:10:10 split for their datasets. This method may not accurately distribute the data and can lead to left or right-skewed MIC values across training, validation, and test splits. To ensure a balanced representation of antimicrobial activity, we split MIC values into quantile-based bins before data partitioning. The number of bins was dynamically determined based on the number of unique MIC measurements to avoid over-fragmentation in smaller datasets. MIC values were grouped to ensure that each bin contained approximately equal numbers of samples. These MIC bins were used solely for stratified data splitting, ensuring proportional representation of low, medium, and high MIC ranges across all splits. This strategy mitigates bias from skewed MIC distributions and enables more reliable evaluation of MIC prediction performance (see Table 1 for more details).

## Methodology

### Protein-Based Language Models in Drug Design

The use of protein-based language models (PLMs), particularly ESM2 [27], marks a significant breakthrough in the computational analysis of protein sequences for drug design. These models effectively leverage deep learning models to create contextual embeddings that capture complex amino acid relationships, essential for predicting the biological activity of potential drug candidates like AMPs. ESM2’s extensive pre-training on protein sequences equips it with valuable representations of evolutionary history and structural characteristics, vital for optimizing therapeutic compounds.

PLMs are transforming AMP research by generating rich, alignment-free sequence representations that outperform traditional descriptors in classification and activity modeling, especially when combined with transfer learning or lightweight predictors [27, 41]. Recent advancements have expanded their applications to complex tasks such as pathogen-specific activity prediction, effectively addressing label imbalance and sparsity [23, 42]. By integrating ESM embeddings with hybrid deep learning frameworks, researchers achieve enhanced performance through comprehensive context and detailed local patterns [43, 44]. Additionally, PLMs are being utilized in generative design pipelines for multi-objective AMP optimization, targeting potency, selectivity, and physicochemical constraints [23, 45]. These developments firmly position ESM-based representations as a cornerstone for next-generation AMP discovery and rational peptide engineering [23, 27, 41, 42, 43, 44, 45].

### ESM-2 Embeddings

ESM2 (Evolutionary Scale Modeling 2) [27] is one of the protein language models (PLMs) available today, serving as a baseline model for protein sequences. It has been trained on an extensive dataset comprised of around 250 million sequences from the UniRef50 database, encompassing a diverse range of organisms, including bacteria, archaea, and eukaryotes. The training process employs masked language modeling (MLM), wherein specific amino acid residues in the input sequences are masked randomly. The model is then tasked with predicting these masked residues based on their contextual relationships within the sequence. This self-supervised training approach enables ESM2 to encode both local and long-range dependencies, yielding contextualized embeddings that implicitly capture functional motifs, biochemical properties, and structural features relevant to the activity of AMPs.

To effectively utilize these embeddings from the ESM2-model, we employ masked mean pooling to aggregate residue-level embeddings into a single and fixed length sequence vector. Since AMP sequences in a batch vary in length, we pad and aggregated the residue level embeddings. This pooled representation serves as a critical input for subsequent predictive tasks, particularly for estimating the MIC of antimicrobial peptides, a key metric for evaluating antimicrobial efficacy.

### Topic Modeling

The integration of motif analysis with topic modeling offers a powerful framework for extracting topic embeddings, especially when paired with ESM2 embeddings for predicting molecular interactions. Motif analysis focuses on identifying significant patterns or subsequences—such as k-mers—that are critical for the biological activity of peptides and other molecular entities. By leveraging topic models, which effectively reveal the contextual relationships among these sequence elements, researchers can better interpret complex biological data. For instance, Schneide et.al proposed a chemical topic modeling [46], using probabilistic framework to categorize large molecular datasets into distinct “chemical topics,” facilitating the discovery of recurring patterns across sets of molecules. Furthermore, PLPTP study [47] incorporated deep learning to enhance the predictive capability for peptide toxicity by capturing evolutionary information while also addressing class imbalance. Ultimately, the synergy between motif-based insights and topic modeling can yield valuable embeddings that enhance the understanding and prediction of molecular behavior, signaling significant advancements in drug development and therapeutic design [48].

To enhance the feature space derived from ESM2 embeddings, we integrate Latent Dirichlet Allocation (LDA), a generative probabilistic model typically used to uncover latent themes in large datasets. Building on our previous research) [26]—which demonstrated that LDA-extracted motifs correlate significantly with biological properties such as MIC, charge, and hydrophobicity—our framework treats protein sequences as “documents” and their constituent overlapping k-mers as “words.” By analyzing the co-occurrence patterns of these k-mers across the entire sequence library, LDA identifies latent topics that represent distinct functional or structural motifs. As shown in Figure 1 A), for any given sequence, the model generates a topic mixture vector that quantifies the prevalence and significance of these motifs. This vector serves as a specialized set of topic embeddings that complements the deep-learning features of ESM2, providing a hybrid representation that captures both global evolutionary context and specific local biochemical patterns. This dual representation strategy—leveraging ESM2 embeddings for localized insights and LDA-derived topic embeddings for broader compositional characteristics, substantially enhances the predictive capacity of our drug design models. The LDA embeddings provide interpretable, low-dimensional summaries of critical biochemical properties, aiding in the identification of sequence patterns that correlate with membrane disruption and antimicrobial effectiveness.

**Fig. 1.**
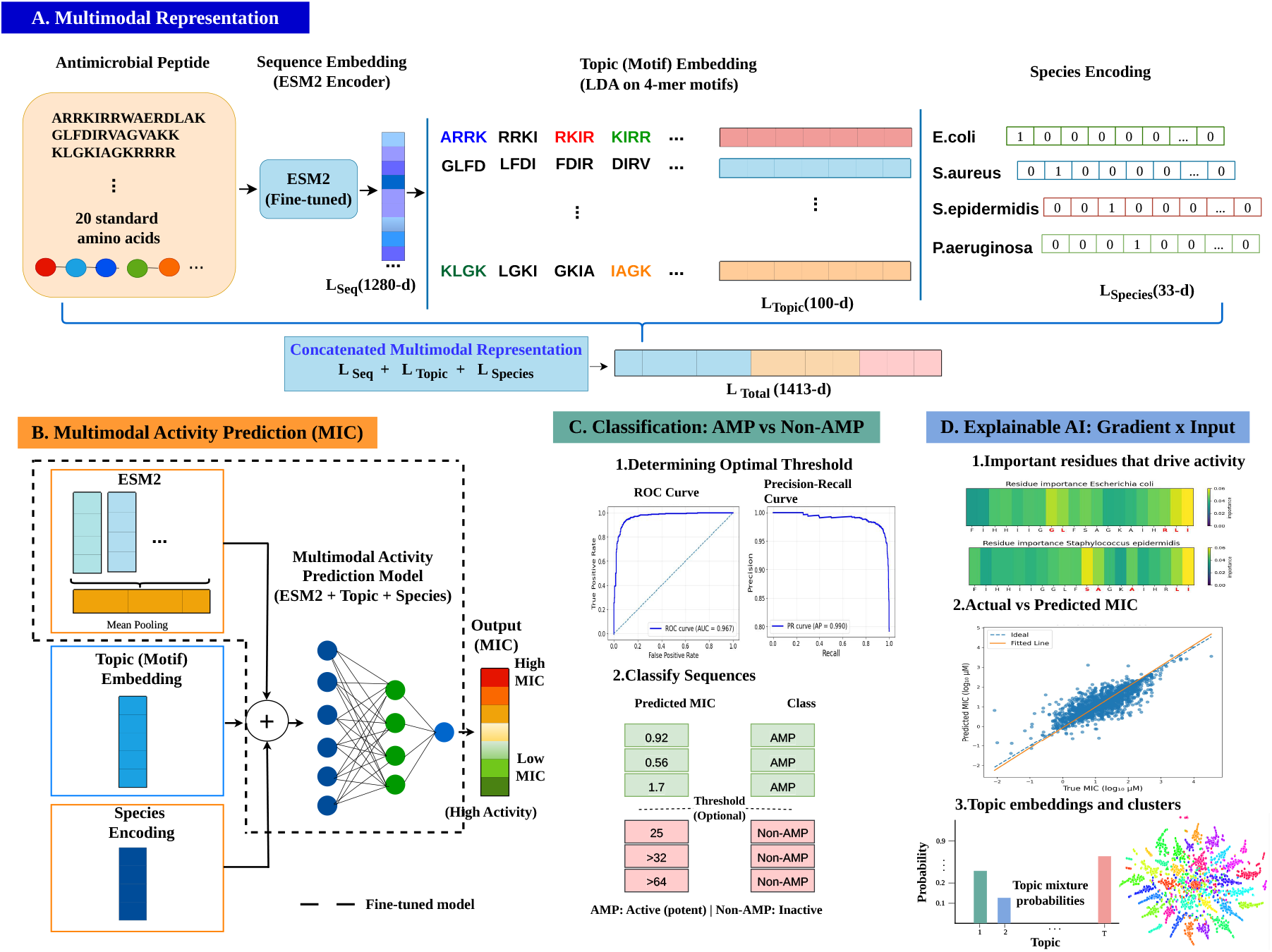
Shows architecture of STAMP (Species and Topic-aware Representation Learning for AMP Discovery). (A) Antimicrobial peptide (AMP) sequences are encoded using ESM2 embeddings. Topic embeddings are derived from an LDA topic model that captures latent antimicrobial motifs, while species one-hot encodings represent the target bacteria. (B) The concatenated representations are fed into a multi-layer perceptron (MLP) regression head to predict Minimum Inhibitory Concentration (MIC) values. The model learns a conditional regression function, represented as *ŷ* = *f*_*θ*_ (*E*(*s*), *T* | *S*), where *E* denotes embeddings, *T* signifies topic embeddings, and *S* represents species encodings. The ESM2 model is fine-tuned for MIC prediction with fixed topic and species encodings. (C) The classification of sequences into AMP and non-AMP categories is achieved using predicted MIC values, with an optimal threshold determined through precision-recall and receiver operating characteristic analyses. (D) An explainable AI component employs the Gradient × Input method to identify key residues that influence antimicrobial activity. Accompanying scatter plots illustrate the correlation between actual and predicted values, and topic embeddings are visualized, showcasing clusters extracted for each sequence via the LDA topic model.

### Species Encoding

In the context of antimicrobial peptide development, it is crucial to account for the variability in MIC across different bacterial species. The same peptide can exhibit very different activities against diverse pathogens due to factors such as variations in membrane composition, cell wall architecture, and resistance mechanisms unique to each species. To address this challenge effectively, we implement a species-aware encoding strategy. Each bacterial species is represented through a categorical encoding, which is then projected into a high-dimensional, dense embedding space.

The benefits of this species-aware encoding are multifaceted. First, it enables the predictive model to condition MIC estimates on the specific susceptibility profiles relevant to the target pathogens, thereby enhancing prediction accuracy. Second, we avoid the pitfalls associated with species-agnostic approaches by incorporating species-specific information directly into the predictive pipeline, allowing for a more realistic representation of peptide-pathogen interactions. This targeted methodology not only enhances the performance of predictive models but also deepens our understanding of the mechanisms by which antimicrobial peptides use their effects across various bacterial species.

### Multimodal Approach

The STAMP model is a multimodal approach for predicting MIC values and classifying AMPs versus non-AMPs. As shown in Figure 1, the model integrates several components: (A) feature embeddings generated from the ESM2 protein language model encapsulate contextualized residue information, while topic embeddings derived from an LDA topic model capture latent motifs relevant to antimicrobial activity. Additionally, species encodings represent specific bacteria through a one-hot representation, contributing to the understanding of target specificity. (B) These diverse embeddings are concatenated to form a unified feature vector, which is then processed through a multi-layer perceptron (MLP) regression head to predict MIC values, allowing the model to learn nonlinear interactions among peptide properties and bacterial targets. (C) The classification of sequences as AMP or non-AMP relies on predicted MIC values, with an optimal decision threshold established through precision-recall and receiver operating characteristic analyses, ensuring reliable differentiation. (D) Furthermore, the model incorporates an explainable AI component using the Gradient × Input method to identify key residues influencing antimicrobial activity, complemented by scatter plots that display the correlation between actual and predicted MIC values. The visualization of topic embeddings and clusters enhances interpretability, linking global compositional trends with specific sequence contexts. This comprehensive framework not only facilitates robust modeling of continuous MIC values but also aligns closely with biological determinants of antimicrobial efficacy, thus advancing the design of effective antimicrobial therapies.

### Problem Formulation

Given a sequence ‘**S**’ and a bacterial species ‘*T* ‘, ‘*y*’ the experimentally measured MIC. The the log-transformed MIC is given as :

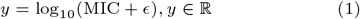

where *ϵ* > 0 is a small constant for numerical stability. The goal is to model the conditional probability of MIC given the peptide sequence and species which is modeled as species aware MIC prediction model:

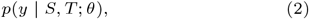

where *θ* represents the model parameters.

Species information is represented using a one-hot encoding projected into a dense embedding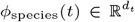. These representations are concatenated and passed through a multilayer perceptron to produce a scalar prediction:

Let the AMP sequence be

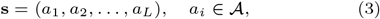

where 𝒜 denotes the amino acid list and *L* is the sequence length. Each sequence *S* is encoded using a pretrained ESM2 protein language model, producing a sequence embedding 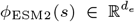. A pretrained ESM2 encoder *f*_ESM_(·) maps the sequence to contextualized residue embeddings:

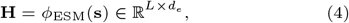

where *d*_*e*_ is the hidden dimension of the ESM2 model.

A fixed-length sequence representation is obtained via pooling:

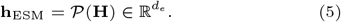

The regression model assumes that the log-transformed MIC follows a Gaussian distribution conditioned on the input features:

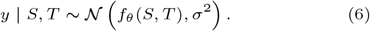

Accordingly, the conditional likelihood is given by:

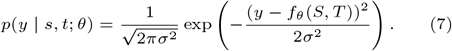

### LDA Topic Mixture Representation

Each AMP sequence is decomposed into overlapping *k*-mers and modeled using Latent Dirichlet Allocation (LDA). For a given sequence, LDA infers a topic mixture vector:

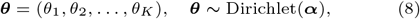

subject to

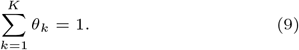

The topic mixture is used directly as a feature vector:

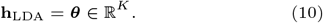

### Species Encoding

Let *S* denote the number of bacterial species. Each species is encoded as a one-hot vector:

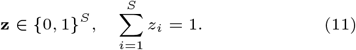

This representation may be projected into a dense embedding:

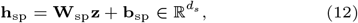

where *d*_*s*_ is the species embedding dimension.

### Regression Model

A multilayer perceptron (MLP) predicts the log-MIC value:

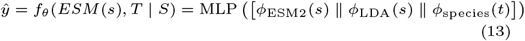

where ∥ denotes vector concatenation, *f*_*θ*_ denotes the nonlinear regression function parameterized by *θ*, ‘x’ is the final feature vector which is constructed by concatenation:

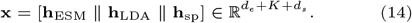

The concatenated feature allows the relationship between sequence features and antimicrobial activity to vary across bacterial targets.

During training, the parameters of both the MLP regression head and the ESM2 encoder are optimized jointly using supervised regression loss, enabling the sequence representation *E*(*s*) to adapt to the MIC prediction task.

### Training Objective

The model is trained by minimizing the mean squared error (MSE) between predicted and observed log-transformed MIC values:

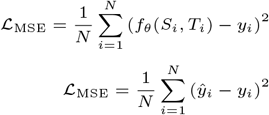

where *N* is the number of training samples. This is equivalent to the maximum likelihood estimation under the Gaussian noise assumption. This formulation integrates contextual sequence embeddings from ESM2, global motif information captured by LDA topic mixtures, and species-specific susceptibility signals into a unified regression framework for MIC prediction.

## Results

We conducted a comprehensive experimental evaluation of the STAMP model’s performance across three benchmark datasets, focusing on its effectiveness in two distinct configurations: constructing species-specific models for individual species and developing a species-aware model that integrates data from multiple species into a singular framework. As detailed in Table 2, our findings indicate that the model consistently achieved a Pearson Correlation Coefficient (PCC) exceeding 0.7, with a score of 0.837 specifically for our curated dataset. This dataset also featured a R-squared (R^2^) score of 0.698 and a low Mean Squared Error (MSE) of 0.171, illustrating the model’s capacity to minimize prediction errors effectively.

**Table 2.** STAMP model evaluated on three benchmark datasets for species- and topic-aware MIC Prediction analysis. Note: Dataset A Refers to benchmark sequences published by Bae et. al [23], Dataset B: refers to benchmark sequences published by Chung et. al [24].

| Dataset | Regression (MIC Prediction) |  |  |  | Classification (AMP vs non-AMP) |  |
| --- | --- | --- | --- | --- | --- | --- |
|  | MSE | MAE | R2 | PCC | Accuracy | Weighted F1 |
| Dataset A (Bae et. al [23]) | 0.275 | 0.385 | 0.531 | 0.732 | 0.87 | 0.88 |
| Dataset B (Chung et. al [24]) | 0.260 | 0.363 | 0.566 | 0.758 | 0.88 | 0.89 |
| Our Curated Dataset | 0.171 | 0.285 | 0.698 | 0.837 | 0.89 | 0.89 |

We compare our proposed multimodal approach for antimicrobial activity prediction with existing state-of-the-art methods. As shown in Table 3, LLAMP achieved performance similar to ours but required building separate models for each of the 35 species and reported only the average score. In contrast, the Ensemble model reported MIC prediction metrics per species by averaging predictions from multiple models. Specifically, they trained eight different models for every species and then selected the top two performing models to obtain average prediction scores. Both the LLAMP and Ensemble models lack generalizability and are computationally expensive. Moreover, our approach boasts superior computational efficiency, allowing for quick adaptations to predict the MIC for unknown sequences.

**Table 3.** Comparison of STAMP method with state-of-the-art methods for MIC prediction analysis. LLAMP♦ built models for each of the 35 species and reported only the average score. Ensemble method⋆ trained eight models for every species and then selected top two performing models to obtain average prediction scores. Our proposed model STAMP is multimodal framework where we train a single model across all species. Note: Models are evaluated by considering the MIC values in logarithmic micromolar (*log*10 (*µM*)) units.

| Dataset | Method | Setting | Species | MSE | MAE | R2 | PCC |
| --- | --- | --- | --- | --- | --- | --- | --- |
| Dataset A | LLAMP [23]♦ | Species-specific | – | 0.272 | 0.379 | 0.536 | 0.735 |
|  | STAMP (Ours) | Species-aware | All species | **0.275** | **0.385** | **0.531** | **0.732** |
| Dataset B | Ensemble model [24]★ | Species-specific | *E.coli*. | 0.225 | – | 0.603 | 0.781 |
|  |  |  | *P.aeruginosa* | 0.20 | – | 0.638 | 0.802 |
|  |  |  | *S.aureus* | 0.274 | – | 0.570 | 0.756 |
|  | STAMP (Ours) | Species-aware | All species | **0.260** | **0.363** | **0.566** | **0.758** |
| Our Dataset | STAMP (Ours) | Species-aware | All species | **0.171** | **0.285** | **0.698** | **0.837** |

Overall, our proposed STAMP model outperforms state-of-the-art models. However, for Dataset B, the Ensemble model achieves a lower MSE for *P. aeruginosa* (0.20) where STAMP achieves MSE of 0.260. Note is that the STAMP’s primary advantage is its generalization and computational efficiency as a single model, not necessarily strictly superior metrics on every single species.

The enhanced performance of our proposed dataset (Our Dataset) can be primarily attributed to through data standardization processes, which may explain the model’s consistent success in predicting MIC values for the *E*.*coli* target across all three datasets. In addition to the PCC of 0.837, the results revealed an *R*^2^ of 0.698, a Mean Absolute Error (MAE) of 0.285, and an MSE of 0.171. These metrics highlight the overall efficacy of the STAMP model in accurately predicting MIC values for AMPs based on key features, including peptide sequences, species encoding, and topic embeddings. This performance significantly surpasses that of baseline models, highlighting the model’s superiority.

To evaluate practical screening performance, the predicted MIC values are categorized into AMP (< 25 *µM*) and non-AMP (> 100*µM*) consistent with the state-of-the-art methods [23]. The resulting confusion matrices reveal strong classification performance across all datasets, with high recall for AMPs and strong specificity for non-AMPs, indicating that the regression predictions maintain biologically meaningful decision boundaries. Overall classification accuracy surpasses 80% across the datasets, reaching as high as 89% with the curated dataset. In practical applications, the MIC predictor is used to prioritize potent candidates by selecting sequences with low predicted MIC values. The observed misclassification of some non-AMPs as AMPs across the datasets is likely due to the limited representation of high-MIC values, which may affect discrimination in the upper MIC range. Despite this, the model consistently demonstrates strong performance across the datasets, supporting its effectiveness for AMP screening and prioritization.

Figures 2, 3, and 4 show scatter plots of predicted versus actual MIC values demonstrating strong agreement across all datasets, with most samples clustering closely around the identity line. This indicates reliable regression performance and the model’s ability to capture the relationships between sequence and activity. We also plotted the receiver operating characteristic (ROC) and precision-recall (PR) curves to automatically determine the threshold for the AMP versus non-AMP classification task. As we notice, ROC and PR are consistently show area under the curve (AUC) and average precision (AP) values exceeding 0.9. This demonstrates robust discrimination between antimicrobial and non-antimicrobial peptides, even in the presence of class imbalance.

**Fig. 2.**
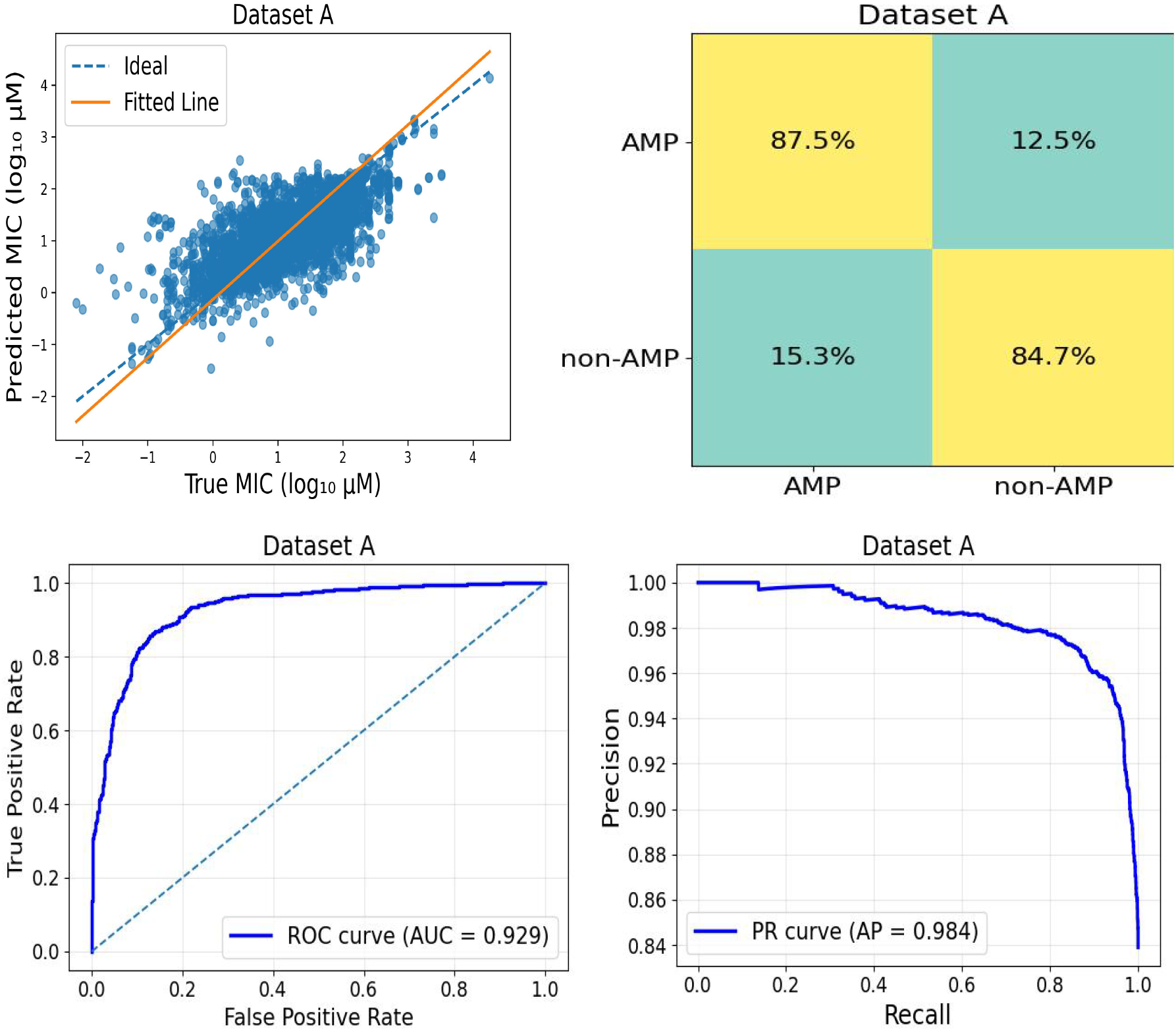
Performance evaluation of MIC prediction model and derived AMP classification on Dataset A.The scatter plot demonstrates strong agreement between predicted and actual MIC values, indicating accurate regression performance. For downstream screening, predicted MIC values were categorized into AMP (<25 *µ* M) and non-AMP (>100 *µ* M) categories. The confusion matrix (%) summarizes classification performance. ROC and precision–recall curves show excellent discriminative capability (AUC(< 0.9, AP (< 0.9), highlighting the model’s robustness and effectiveness in identifying antimicrobial peptides under class imbalance.

**Fig. 3.**
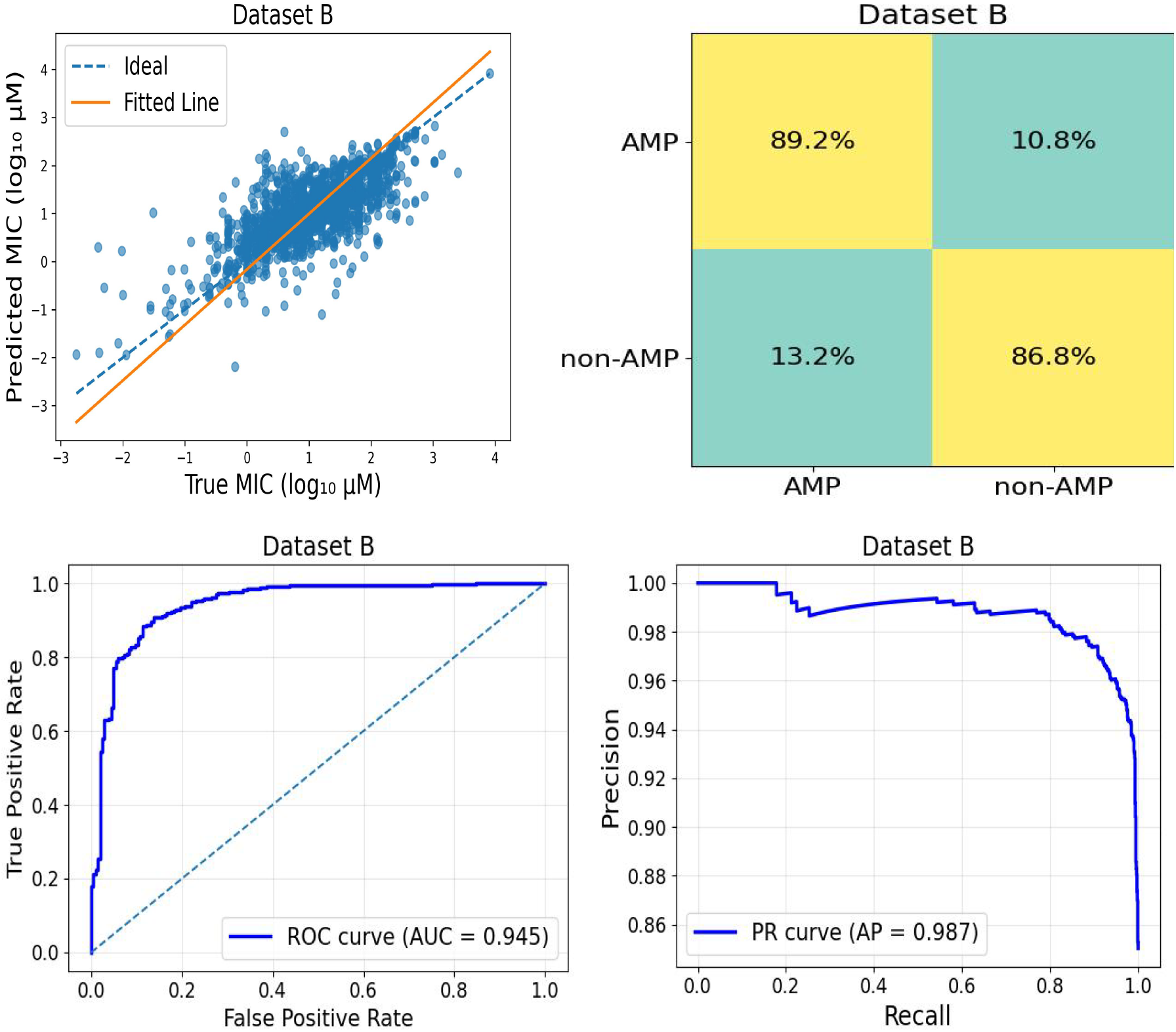
Performance evaluation of MIC prediction model and derived AMP classification on Dataset B. Note: full description is provided in Figure 2.

**Fig. 4.**
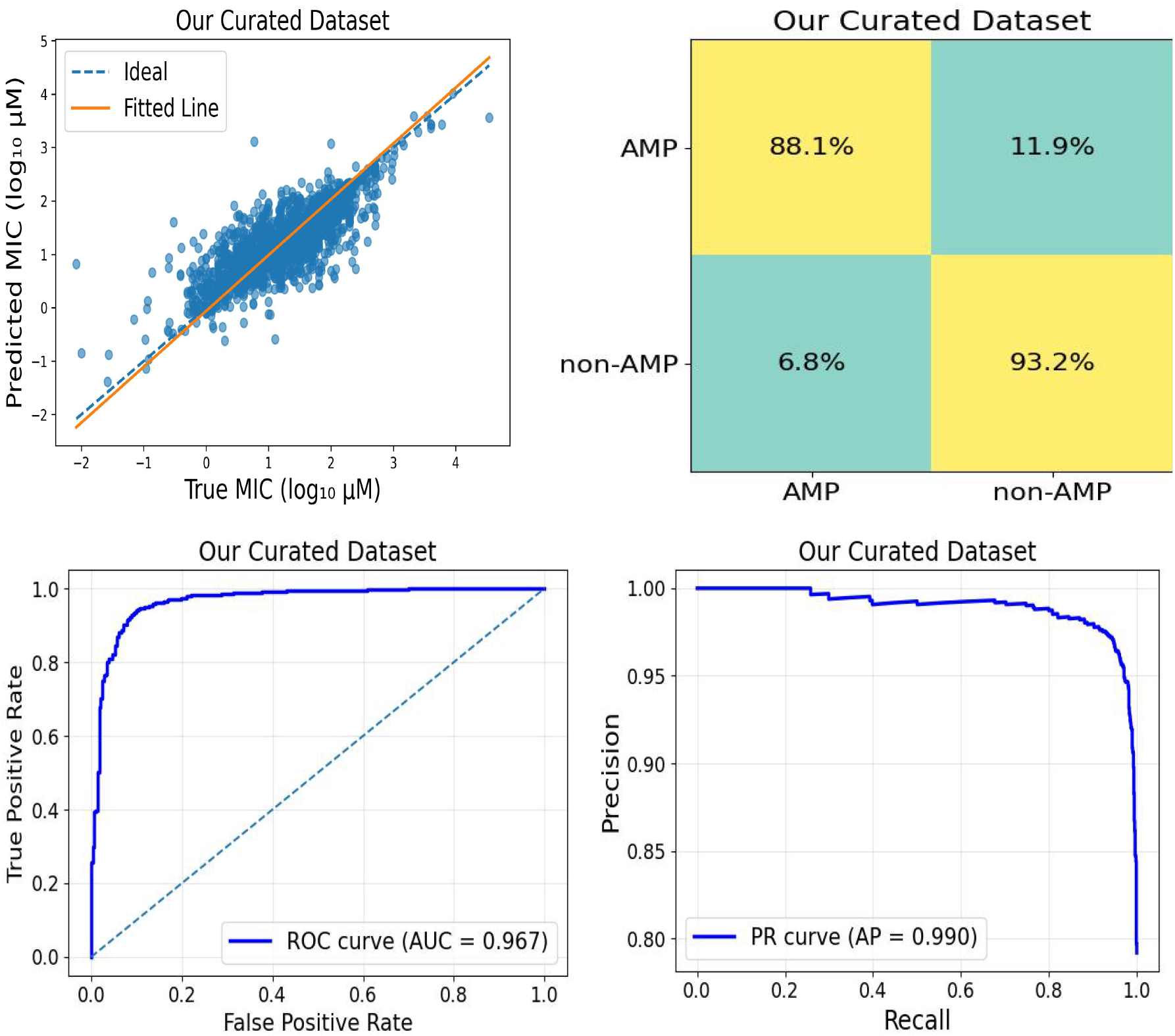
Performance evaluation of MIC prediction model and derived AMP classification on our curated dataset. Note: full description is provided in Figure 2.

We further assessed the proposed STAMP model based on a species-specific across the three datasets. Tables S1, S2, and S3 in Supplementary Material present a detailed summary of the model’s species-specific performance. For the Dataset A and our scraped dataset, the model consistently demonstrated a PCC score exceeding 0.7 and *R*^2^ values above 0.5. Notably, the model showed good performance for the *E*.*coli* target, outperforming other species in the comparison. Overall, the model achieved consistent results across all targets in our curated dataset, with particular strength in predicting values for *E*.*coli*.

We further performed bias analysis of STAMP model predictions across the three datasets. As shown in Table 4, prediction bias analysis of three datasets indicate that the model does not exhibit systematic over- or underestimation of MIC values. Specifically, the mean over-prediction and under-prediction is comparable across the datasets and the errors are distributed relatively symmetrically around the true MIC values. While the Dataset B showed slightly higher variance in its error margins, the overall lack of a dominant directional skew confirms that the model’s predictive logic remains stable across diverse protein sequences and MIC ranges.

**Table 4.** Prediction bias and regression stability analysis across datasets. The mean over-prediction and under-prediction is comparable across the datasets and the errors are distributed relatively symmetrically around the true MIC values indicating that the model does not exhibit systematic over- or underestimation of MIC values

| Dataset | Over Predictions ($\mu \pm \sigma$) | Under Predictions ($\mu \pm \sigma$) |
| --- | --- | --- |
| Dataset A | $0.373 \pm 0.312$ | $-0.279 \pm 0.278$ |
| Dataset B | $0.373 \pm 0.377$ | $-0.351 \pm 0.356$ |
| Our Curated Dataset | $0.310 \pm 0.312$ | $-0.279 \pm 0.278$ |

### Species-Aware Feature Learning

We further validate the performance of STAMP by predicting MIC values for peptides experimentally tested against two bacterial strains: the Gram-positive *S*.*epidermidis* and the Gram-negative *E*.*coli*. In our earlier study [28], we investigated the role of C-terminal poly-arginine segments in enhancing antimicrobial activity using the fish-derived peptide Tilapia piscidin 4 (TP4) and its truncated variant lacking the poly-arginine tail (TP4-noR5). We also examined a neutral peptide (NP) with no intrinsic antimicrobial activity; however, appending a poly-arginine tail to its C-terminus significantly improved its efficacy. Table 5 shows the peptides and corresponding MIC measurements against both pathogens. For further details on MIC measurements and assay protocols, please refer to our earlier study [28].

**Table 5.**
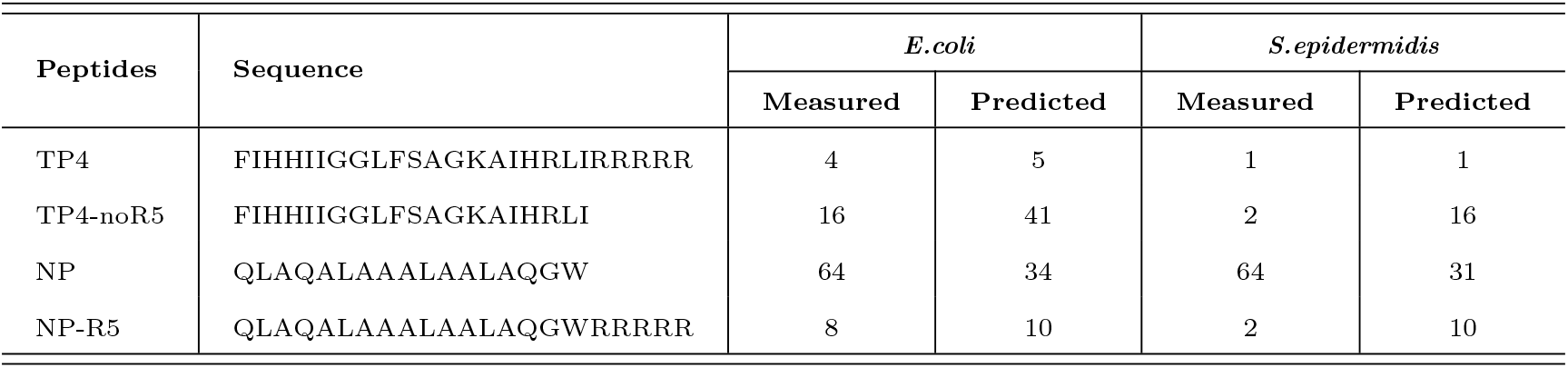
STAMP model MIC predictions vs measured MIC values for TP4 sequences. Our proposed model misclassify one sequence (TP4-noR5) as non-AMP against E.coli. The STAMP model demonstrates target-dependent predictions while preserving consistent activity trends. The two sequences enriched with C-terminal arginine clusters (TP4 and NP-R5) exhibit consistent predicted MIC values across both *E*.*coli* and *S*.*epidermidis*, closely aligning with Kaur et.al [28] experimentally determined low MIC values. We can also notice that the difference between actual and predicted MICs for these sequences is consistent across targets. Note: MIC values are reported in *µM*.

Table 5 shows that for the four TP4-derived peptides, the model demonstrates target-dependent predictions while preserving consistent activity trends. The two sequences enriched with C-terminal arginine clusters (TP4 and NP-R5) exhibit consistent predicted MIC values across both *E*.*coli* and *S*.*epidermidis*, closely aligning with their experimentally low MIC values. We can also notice that the difference between actual and predicted MICs for these sequences is consistent across targets. This suggests that the model reliably recognizes strongly cationic, arginine-rich motifs as broadly active across Gram-negative and Gram-positive bacteria.

On the other hand, the other two sequences lacking extended C-terminal arginine tails show pronounced target-dependent shifts in predicted activity. For example, TP4-noR5 displays different experimental MICs between *E*.*coli* (16 *µ*M) and *S. epidermidis* (2 *µ*M), and the model correspondingly adjusts its predictions (41 *µ*M vs 16 *µ*M). Similarly, neutral peptide (NP) shows moderate predictive variation between targets while maintaining its overall weak activity prediction. These target-specific prediction shifts indicate that the model does not apply a uniform sequence-to-activity mapping, but instead modulates its predictions based on species context.

To further understand which amino acids matter most for predicting antimicrobial activity, we created heatmaps that show residues importance for AMP activity. To compute residue importance for AMP activity, we utilized a Gradient × Input saliency approach [49] to map MIC sensitivity back to individual amino acids. By calculating the gradients of the predicted MIC with respect to the ESM2 hidden states and multiplying them by the original embeddings, we quantified the contribution of each residue to the final output. These raw importance scores were masked to remove padding and special tokens (`<cls>`, `<eos>`), then normalized across the sequence to ensure comparability. The resulting saliency values were visualized as heatmaps, allowing for the direct identification of specific residues and structural motifs that play key role in predicting MIC of a given AMPs.

To analyze residue importance, we focused on TP4 sequences, using data where the same sequences have MIC measurements and predictions for both *E*.*coli* and *S*.*epidermidis*. For identical TP4-derived sequences (a & b, c & d, e & f, g & h), the heatmaps reveal distinct patterns of amino acid contributions when conditioned on different bacterial species. Specifically, residues that are highly influential for MIC prediction in *E*.*coli* are not necessarily the same residues emphasized under *S*.*epidermidis* conditioning. These species-dependent shifts in residue importance indicate that the model does not rely on a static sequence representation, but instead dynamically re-weights sequence features depending on the bacterial species. This suggests that species context modulates how the model interprets peptide physicochemical properties, such as charge distribution, hydrophobicity, and amphipathicity, which are known to differentially influence membrane interactions in Gram-negative and Gram-positive bacteria. The divergence in attribution patterns provides mechanistic evidence that the model has learned species-aware antimicrobial susceptibility signatures at the residue level.

Figure 5 shows, top five important amino acids are emphasized (red in color) for MIC prediction analysis. The analysis of residue importance of positively charged residues, particularly arginine, among the most influential positions for MIC prediction. These residues likely contribute to electrostatic interactions with negatively charged bacterial membranes. As illustrated in Figures 5 a, g, and h, the poly-arginine (poly-R) motifs located at the C-terminus exhibit significant importance for AMP activity. This finding aligns with the experimental results reported by Kaur et.al [28], which demonstrate that poly-arginine tails enhance antimicrobial characteristics within a sequence. These heatmaps suggest that the model has correctly identified these cationic residues as critical structural determinants for the predicted MIC values. Hydrophobic residues such as leucine(L), isoleucine(I), alanine(A), and tryptophan(W) were also frequently identified, suggesting that the model captures amphipathic sequence patterns associated with membrane insertion and disruption. In addition, glycine (G) residues appeared among important positions, potentially reflecting their role in introducing structural flexibility or hinge regions within antimicrobial peptides. Differences in residue importance between *E*.*coli* and *S*.*epidermidis* targets indicate that the model learns species-specific sequence features relevant to antimicrobial activity.

**Fig. 5.**
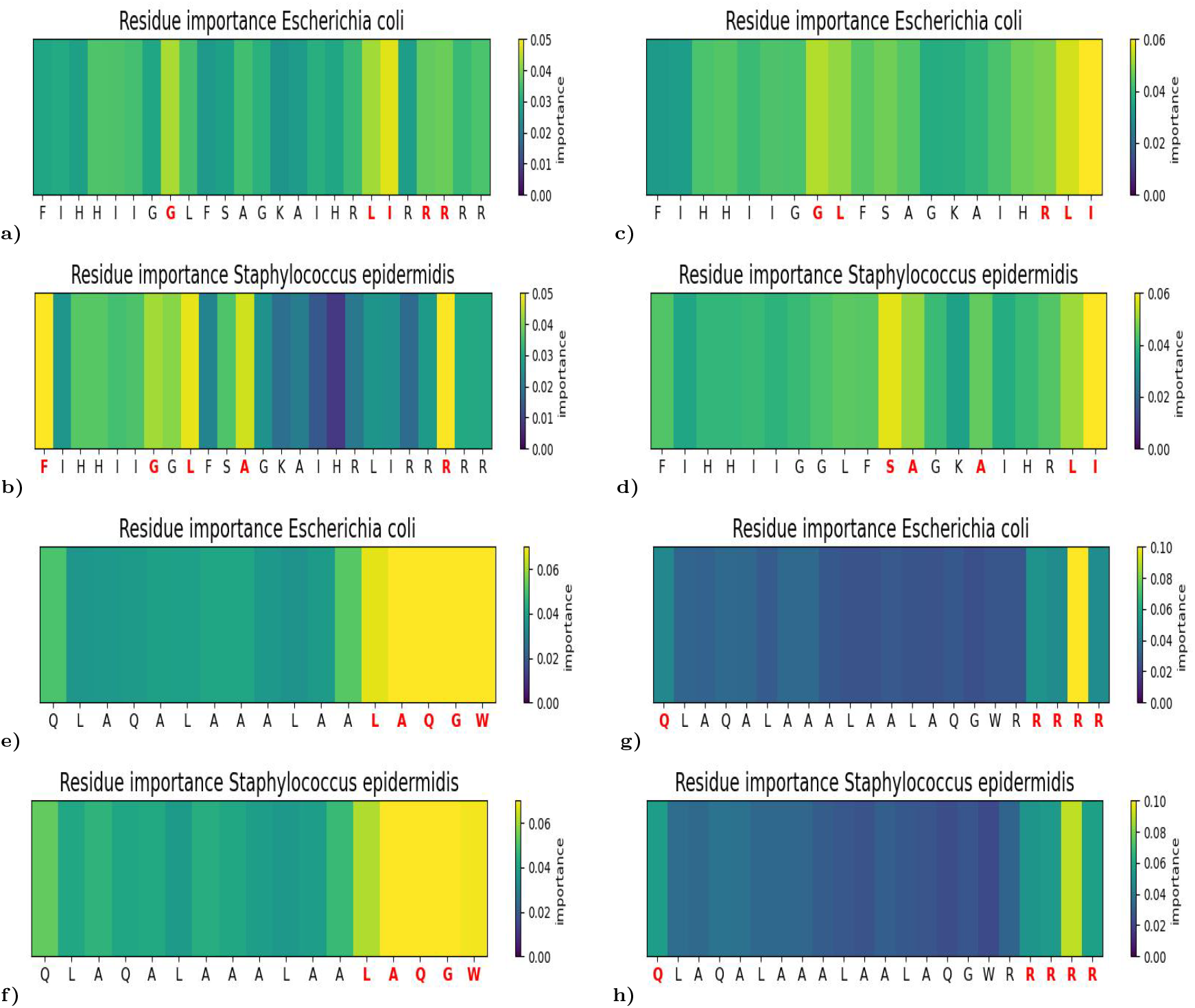
Residue importance analysis shows enrichment of positively charged residues, particularly arginine, among the most influential positions for MIC prediction. Hydrophobic residues such as leucine, isoleucine, and tryptophan are also frequently highlighted, suggesting that the model captures amphipathic sequence features associated with antimicrobial peptide membrane disruption. Differences in residue importance between *E*.*coli* and *S*.*epidermidis* targets further indicate species-dependent sequence feature recognition by the model.

## Ablation Experiments

We conducted a rigorous comparative analysis of our STAMP model against the methodology outlined by Chung et al. [24]. Table 3 details the performance metrics from the dataset proposed by Chung et al. Our evaluation of a single model across three target species yielded a PCC of 0.758 and an R^2^ score of 0.566. While Chung et al. reported superior performance metrics, their methodology involved constructing eight distinct models for each target species, leading to a total of 24 models. Afterward, they selected the top two models for each target, which varied significantly in effectiveness depending on the specific target.

It is important to note that the models developed by Bae et al. [23] are heavily reliant on ESM2 embeddings and genomic features that may not translate effectively to other bacterial strains beyond those explicitly mentioned in their study. In stark contrast, our proposed STAMP model adeptly harnesses ESM2 embeddings in conjunction with topic mixtures to capture nuanced categories based on k-mers and species encoding.

Table 3 shows that, our model demonstrates good performance for *E*.*coli* and *S. aureus* targets, despite some notable errors when predicting for *P*.*aeruginosa*. Nevertheless, the overall performance of our single model achieves a PCC of 0.758 and a low MSE of 0.260. To comprehensively assess the impact of finetuning the ESM2 model, along with incorporating LDA topic model embeddings and species encoding, we conducted a rigorous evaluation of several target-specific models. Our analyses involved multiple configurations, including ESM2 embeddings combined with a regression layer specifically designed for MIC prediction, ESM2 embeddings further augmented with topic embeddings followed by regression, and the strategic finetuning of the ESM2 model using LDA topic embeddings. The results, as clearly illustrated in Tables S1, S2, and S3 in supplementary material (models are shown in Figure S1), reveal that the fine-tuned ESM2 model utilizing LDA topic embeddings outperformed all baseline models across three benchmark datasets and a diverse range of target species. This compelling evidence demonstrates that the model effectively learns and captures species-specific features through the meticulous finetuning of ESM2 in conjunction with topic embeddings.

Moreover, we examined the performance of each ESM2 embedding model in terms of their architecture, which could consist of 8 million (8M), 35 million (35M), or 650 million (650M) parameters for extracting embeddings and subsequent finetuning. By analyzing the model performances associated with these three distinct ESM configurations (Tables S1, S2, and S3 in supplementary material), we found that the 8M parameter ESM2 model excelled in finetuning scenarios, outperforming both the 35M and 650M models. Conversely, for tasks focused on feature extraction, the 650M model demonstrated significant advantages, outperforming the 8M and 35M models. This indicates that finetuning of 8M ESM2 model requires substantially less training data compared to the larger models, making it potentially more accessible for scenarios with limited data availability. On the other hand, the embeddings generated by the 650M ESM2 model were exceptionally robust, demonstrating a notable efficacy in MIC prediction analysis, thereby highlighting its capacity to handle complex prediction tasks with high reliability.

## Conclusion

This study presents STAMP (Species- and Topic-aware Representation Learning for Antimicrobial Peptide Discovery), a novel multimodal framework for cross-species antimicrobial activity prediction that leverages protein-based Large Language Model (ESM-2) along with species- and topic-aware embeddings for unseen sequences based on specific pathogens. Our findings indicate that STAMP effectively generalizes across species-specific features, capturing key biological determinants of antimicrobial activity. We validated STAMP’s robustness using three independent datasets, achieving a Pearson Correlation Coefficient 0.837 and an*R*^2^ of 0.7, with high ROC and precision-recall values, both exceeding 0.9. These results confirm our proposed model’s robustness in predicting the MIC of specific peptides and categorizing them as antimicrobial or non-antimicrobial, facilitating the identification of peptides that could serve as potential AMPs for drug design analysis. Additionally, we analyzed residue-level importance using the gradient × Input method to uncover critical sequence determinants for antimicrobial activity. In future work, we plan to integrate genomic features and physicochemical descriptors to enhance predictive accuracy and continue to standardize antimicrobial databases for the research community.

## Competing interests

The authors declare no competing interests.

## Author contributions statement

SP, AC, DH, FH, MM, JKB and AC conceived the study. SP developed the methodology, was responsible for the computational evaluations and carried out formal analysis. SP and KM curated data, NK measured MICs for TP4 sequences. SP wrote the original article draft, which was reviewed by DH, FH, MM, JBK and AC. SP was responsible for visualizations. MM, FH, DH, JBK, and AC supervised this study. WK contributed to the critical revision of the manuscript for important intellectual content.

## Funding

This work was supported by an Innovation in Measurement Science (IMS) grant from the National Institute of Standards and Technology (NIST) (70NANB17H299 and 70NANB24H248).

## Disclaimer

The commercial products used in this study were only referenced to specify the experimental procedure adequately. Such identification of commercial products is not intended to imply recommendation or endorsement by the National Institute of Standards and Technology, nor is it intended to imply that the identified products are necessarily the best available for the purpose.

## Notes

### Competing Interest Statement

The authors have declared no competing interest.

