## supplementary file for "Species- and Topic-aware Representation Learning for Antimicrobial Peptide Discovery"

### 1 Metrics

To evaluate the performance of our proposed STAMP model for the MIC prediction task, we use both error-based and correlation-based metrics to provide a comprehensive assessment. For error-based metrics, we employ Mean Squared Error (MSE) and Mean Absolute Error (MAE). These metrics quantify the deviation between predicted and experimentally measured MIC values, reflecting the absolute accuracy of the predictions in biologically meaningful units (or their log-transformed equivalents). MAE is particularly interpretable, as it indicates the typical magnitude of prediction errors without disproportionately penalizing outliers. In contrast, MSE emphasizes larger errors and is sensitive to extreme mispredictions, which may be critical for assessing antimicrobial efficacy.

On the other hand, Pearson correlation coefficient (PCC) and the coefficient of determination ($R^2$) primarily evaluate the strength of linear association and the variance explained by the model, respectively. However, they do not account for systematic prediction bias or the magnitude of absolute errors. A model may demonstrate a high correlation while still generating clinically unacceptable MIC estimates if the predictions are consistently shifted or scaled. Therefore, it is crucial to report MSE and MAE alongside correlation-based metrics to accurately characterize predictive performance and ensure practical relevance in antimicrobial peptide MIC prediction tasks.

#### 1.1 The Coefficient of Determination ($R^2$)

The coefficient of determination ($R^2$) is a metric that measures how well the independent variable (x) explains the variation in the dependent variable (Y) within a regression model. When calculating $R^2$ between true values and predicted values, we can assess the model's performance. A higher $R^2$ value indicates that the model explains a significant portion of the variability in true values, while a low $R^2$ suggests that the model fails to capture much variance and may be overlooking key data patterns. If $R^2$ is negative, it indicates that the model performs worse than simply predicting the mean value of y. The formula for $R^2$ when comparing actual values to predicted values is as follows:

$$R^2 = 1 - \frac{\sum(y_{\text{true}} - y_{\text{pred}})^2}{\sum(y_{\text{true}} - \bar{y})^2}$$

where: $y_{\text{true}}$ = actual values
$y_{\text{pred}}$ = model predictions
$\bar{y}$ = mean of actual values
If $R^2 = 1$, the model predicts perfectly.
If $R^2 = 0$, the model is no better than predicting the mean of y.
If $R^2 < 0$, the model is worse than a naive guess. (1)

#### 1.2 Pearson Correlation Coefficient (PCC):

The Pearson correlation coefficient (PCC) is commonly used to measure how well predicted values correlate with actual values in regression analysis. It assesses the alignment of predicted values with actual values but does not provide information about the variance explained by the model. PCC measures the linear relationship between actual values ($y_{\text{true}}$) and predicted values ($y_{\text{pred}}$), indicating the strength and direction (positive or negative) of the correlation between the two variables. It is computed as:

$$r = \frac{\sum(y_{\text{true}} - \bar{y})(y_{\text{pred}} - \bar{y}_{\text{pred}})}{\sqrt{\sum(y_{\text{true}} - \bar{y})^2} \cdot \sqrt{\sum(y_{\text{pred}} - \bar{y}_{\text{pred}})^2}}$$

where $r = 1$ means perfect positive correlation, $r = -1$ indicates a negative correlation, and $r = 0$ means no correlation. (2)

### 2 Ablation Experiments

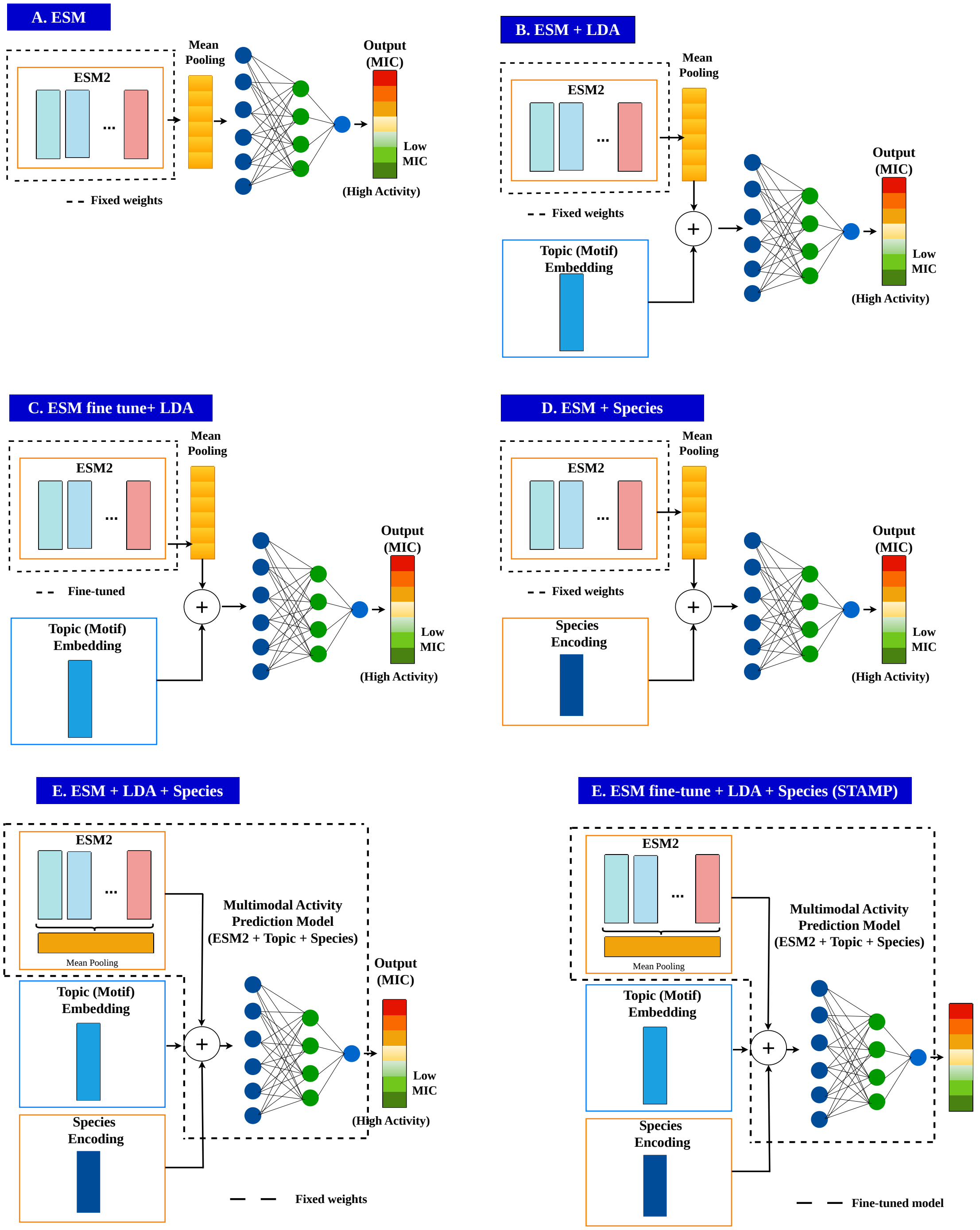


Figure S1: Illustrates the various model architectures evaluated during the ablation experiments, which were designed to compare the performance in predicting AMP activity as detailed in Tables S1, S2, S3 A. The ESM model is employed to extract embeddings from sequences, with a regression model developed on top of these fixed embeddings. B. This architecture is similar to A but incorporates topic embeddings to enhance the feature vectors from the ESM model. C. In addition to the ESM model and topic embeddings, this architecture includes a regression head and fine-tunes the ESM for AMP activity prediction. D. This architecture concatenates ESM embeddings with species encodings before passing them to the regression model. E. This approach combines ESM embeddings, topic embeddings, and species encodings and feeds them into a regression layer for predicting AMP activity. F. In this final architecture, ESM embeddings, topic embeddings, and species encodings are concatenated, and the regression model along with the ESM model is fine-tuned end-to-end for AMP activity prediction (STAMP). This comprehensive overview highlights the different strategies employed to leverage ESM embeddings and additional features for optimizing AMP activity prediction.

Table S1: Ablation study comparing ESM2 pretrained models (8M, 35M and 650M parameter models) for MIC prediction analysis for Dataset A. Performance is evaluated using regression (MAE, PCC, $\mathbf{R^2}$), with the best results in each column highlighted in bold.

| **Dataset** | **Species** | **MSE** | **MAE** | $R^2$ | **PCC** | **Model** |
|---|---|---|---|---|---|---|
| ESM | E.coli | 0.301 | 0.413 | 0.491 | 0.701 | 8M |
| | | 0.295 | 0.407 | 0.501 | 0.712 | 35M |
| | | 0.297 | 0.414 | 0.498 | 0.711 | 650M |
| | P.aeruginosa | 0.280 | 0.404 | 0.465 | 0.682 | 8M |
| | | 0.282 | 0.40 | 0.462 | 0.681 | 35M |
| | | 0.279 | 0.390 | 0.467 | 0.691 | 560M |
| | S. aureus | 0.347 | 0.462 | 0.384 | 0.621 | 8M |
| | | 0.313 | 0.430 | 0.445 | 0.674 | 35M |
| | | 0.322 | 0.429 | 0.429 | 0.659 | 650M |
| | S. Epidermidis | 0.277 | 0.385 | 0.441 | 0.678 | 8M |
| | | 0.279 | 0.393 | 0.437 | 0.679 | 35M |
| | | 0.286 | 0.385 | 0.424 | 0.669 | 650M |
| ESM + LDA | E.coli | 0.294 | 0.406 | 0.503 | 0.714 | 8M |
| | | 0.276 | 0.387 | 0.525 | 0.732 | 35M |
| | | 0.281 | 0.397 | 0.524 | 0.726 | 650M |
| | P.aeruginosa | 0.256 | 0.382 | 0.511 | 0.719 | 8M |
| | | 0.269 | 0.389 | 0.487 | 0.702 | 35M |
| | | 0.277 | 0.394 | 0.470 | 0.691 | 650M |
| | S. aureus | 0.323 | 0.446 | 0.426 | 0.653 | 8M |
| | | 0.299 | 0.418 | 0.469 | 0.690 | 35M |
| | | 0.325 | 0.432 | 0.423 | 0.654 | 650M |
| | S. Epidermidis | 0.255 | 0.369 | 0.487 | 0.705 | 8M |
| | | 0.261 | 0.379 | 0.474 | 0.697 | 35M |
| | | 0.269 | 0.368 | 0.457 | 0.694 | 650M |
| ESM Finetune + LDA | E.coli | 0.252 | 0.364 | 0.574 | 0.763 | 8M |
| | | 0.248 | 0.363 | 0.580 | 0.763 | 35M |
| | | **0.240** | **0.357** | **0.595** | **0.771** | 650M |
| | P.aeruginosa | 0.238 | 0.348 | 0.546 | 0.744 | 8M |
| | | **0.225** | **0.338** | **0.570** | **0.756** | 35M |
| | | 0.233 | 0.349 | 0.555 | 0.747 | 650M |
| | S. aureus | 0.281 | 0.386 | 0.5 | 0.713 | 8M |
| | | **0.266** | **0.378** | **0.527** | **0.728** | 35M |
| | | 0.285 | 0.389 | 0.494 | 0.704 | 650M |
| | S. Epidermidis | 0.213 | 0.335 | 0.571 | 0.76 | 8M |
| | | 0.239 | 0.351 | 0.562 | 0.729 | 35M |
| | | **0.217** | **0.337** | **0.562** | **0.760** | 650M |
| ESM +Species | Species-aware | 0.390 | 0.490 | 0.334 | 0.581 | 8M |
| | | 0.386 | 0.479 | 0.341 | 0.589 | 35M |
| | | 0.40 | 0.491 | 0.318 | 0.570 | 560M |
| ESM+LDA + Species | Species-aware | 0.292 | 0.399 | 0.502 | 0.717 | 8M |
| | | 0.299 | 0.405 | 0.490 | 0.706 | 35M |
| | | 0.311 | 0.421 | 0.470 | 0.687 | 650M |
| ESM-Finetune + LDA + Species (STAMP) | Species-aware | **0.275** | **0.385** | **0.531** | **0.732** | 8M |
| | | 0.280 | 0.391 | 0.522 | 0.729 | 35M |
| | | 0.295 | 0.405 | 0.496 | 0.706 | 650M |

Table S2: Ablation study comparing ESM2 pretrained models (8M, 35M and 650M parameter models) for MIC prediction analysis for Dataset B. Performance is evaluated using regression (MAE, PCC, $\mathbf{R^2}$), with the best results in each column highlighted in bold.

| **Dataset** | **Species** | **MSE** | **MAE** | $R^2$ | **PCC** | **Model** |
|---|---|---|---|---|---|---|
| ESM | E.coli | 0.307 | 0.422 | 0.457 | 0.687 | 8M |
| | | 0.309 | 0.419 | 0.453 | 0.682 | 35M |
| | | 0.302 | 0.418 | 0.467 | 0.688 | 650M |
| | P.aeruginosa | 0.368 | 0.447 | 0.352 | 0.610 | 8M |
| | | 0.372 | 0.457 | 0.346 | 0.602 | 35M |
| | | 0.368 | 0.439 | 0.351 | 0.604 | 650M |
| | S. aureus | 0.406 | 0.485 | 0.363 | 0.608 | 8M |
| | | 0.410 | 0.490 | 0.357 | 0.606 | 35M |
| | | 0.424 | 0.502 | 0.335 | 0.597 | 650M |
| ESM + LDA | E.coli | 0.302 | 0.427 | 0.466 | 0.694 | 8M |
| | | 0.296 | 0.417 | 0.476 | 0.695 | 35M |
| | | 0.298 | 0.412 | 0.473 | 0.699 | 650M |
| | P.aeruginosa | 0.348 | 0.428 | 0.387 | 0.638 | 8M |
| | | 0.360 | 0.447 | 0.365 | 0.611 | 35M |
| | | 0.358 | 0.435 | 0.369 | 0.624 | 650M |
| | S. aureus | 0.372 | 0.466 | 0.416 | 0.653 | 8M |
| | | 0.388 | 0.485 | 0.390 | 0.633 | 35M |
| | | 0.407 | 0.487 | 0.361 | 0.616 | 650M |
| ESM finetune + LDA | E.coli | 0.258 | 0.375 | 0.544 | 0.745 | 8M |
| | | **0.255** | **0.378** | **0.549** | **0.746** | 35M |
| | | 0.274 | 0.384 | 0.515 | 0.728 | 650M |
| | P.aeruginosa | **0.337** | **0.410** | **0.407** | **0.659** | 8M |
| | | 0.363 | 0.419 | 0.361 | 0.637 | 35M |
| | | 0.338 | 0.410 | 0.405 | 0.657 | 650M |
| | S. aureus | 0.350 | 0.443 | 0.451 | 0.687 | 8M |
| | | **0.321** | **0.422** | **0.497** | **0.714** | 35M |
| | | 0.322 | 0.433 | 0.494 | 0.706 | 650M |
| ESM + Species | Species-aware | 0.347 | 0.446 | 0.421 | 0.654 | 8M |
| | | 0.346 | 0.451 | 0.422 | 0.660 | 35M |
| | | 0.336 | 0.440 | 0.438 | 0.668 | 650M |
| ESM + LDA +Species | Species-aware | 0.291 | 0.402 | 0.513 | 0.721 | 8M |
| | | 0.280 | 0.392 | 0.532 | 0.740 | 35M |
| | | 0.271 | 0.387 | 0.547 | 0.748 | 650M |
| ESM-Finetune + LDA + Species (STAMP) | Species-aware | 0.321 | 0.431 | 0.464 | 0.682 | 8M |
| | | **0.260** | **0.363** | **0.566** | **0.758** | 35M |
| | | 0.319 | 0.419 | 0.467 | 0.701 | 650M |

Table S3: Ablation study comparing ESM2 pretrained models (8M, 35M and 650M parameter models) for MIC prediction analysis for Our Curated Dataset. Performance is evaluated using regression (MAE, MSE, PCC, $\mathbf{R^2}$), with the best results in each column highlighted in bold.

| **Dataset** | **species** | **MSE** | **MAE** | $R^2$ | **PCC** | **Model** |
|---|---|---|---|---|---|---|
| ESM | E.coli | 0.296 | 0.404 | 0.521 | 0.723 | 8M |
| | | 0.293 | 0.409 | 0.526 | 0.728 | 35M |
| | | 0.299 | 0.408 | 0.516 | 0.725 | |
| | P.aeruginosa | 0.288 | 0.470 | 0.405 | 0.636 | 8M |
| | | 0.278 | 0.405 | 0.425 | 0.625 | 35M |
| | | 0.269 | 0.402 | 0.443 | 0.666 | 650M |
| | S. aureus | 0.307 | 0.433 | 0.425 | 0.655 | 8M |
| | | 0.293 | 0.420 | 0.451 | 0.678 | 35M |
| | | 0.307 | 0.435 | 0.425 | 0.655 | 650M |
| | S. Epidermidis | 0.269 | 0.412 | 0.534 | 0.735 | 8M |
| | | 0.247 | 0.375 | 0.572 | 0.758 | 35M |
| | | 0.278 | 0.394 | 0.519 | 0.732 | 560M |
| ESM + LDA | E.coli | 0.277 | 0.390 | 0.552 | 0.749 | 8M |
| | | 0.277 | 0.394 | 0.552 | 0.746 | 35M |
| | | 0.264 | 0.382 | 0.573 | 0.763 | 650M |
| | P.aeruginosa | 0.250 | 0.377 | 0.482 | 0.698 | 8M |
| | | 0.273 | 0.405 | 0.436 | 0.663 | 35M |
| | | 0.261 | 0.393 | 0.460 | 0.681 | 650M |
| | S. aureus | 0.270 | 0.405 | 0.494 | 0.710 | 8M |
| | | 0.280 | 0.417 | 0.475 | 0.695 | 35M |
| | | 0.269 | 0.403 | 0.497 | 0.714 | 650M |
| | S. Epidermidis | 0.257 | 0.393 | 0.555 | 0.746 | 8M |
| | | 0.268 | 0.391 | 0.535 | 0.735 | 35M |
| | | 0.271 | 0.389 | 0.531 | 0.741 | 650M |
| ESM finetune + LDA | E.coli | 0.258 | 0.363 | 0.582 | 0.770 | 8M |
| | | 0.263 | 0.359 | 0.574 | 0.762 | 35M |
| | | **0.247** | **0.355** | **0.60** | **0.775** | 650M |
| | P.aeruginosa | 0.216 | 0.334 | 0.552 | 0.751 | 8M |
| | | 0.212 | 0.335 | 0.561 | 0.753 | 35M |
| | | **0.209** | **0.336** | **0.568** | **0.756** | 650M |
| | S. aureus | 0.250 | 0.369 | 0.532 | 0.738 | 8M |
| | | 0.243 | 0.369 | 0.544 | 0.744 | 35M |
| | | **0.232** | **0.367** | **0.565** | **0.753** | 650M |
| | S. Epidermidis | **0.236** | **0.351** | **0.590** | **0.770** | 8M |
| | | 0.254 | 0.367 | 0.560 | 0.749 | 32M |
| | | 0.253 | 0.360 | 0.561 | 0.754 | 650M |
| ESM +Species | Species-aware | 0.264 | 0.399 | 0.532 | 0.730 | 8M |
| | | 0.266 | 0.401 | 0.529 | 0.727 | 35M |
| | | 0.261 | 0.389 | 0.538 | 0.736 | 650M |
| ESM + LDA + Species | Species-aware | 0.211 | 0.343 | 0.626 | 0.794 | 8M |
| | | 0.210 | 0.345 | 0.628 | 0.794 | 35M |
| | | 0.20 | 0.337 | 0.645 | 0.805 | 650M |
| ESM-Finetune + LDA + Species (STAMP) | Species-aware | **0.171** | **0.285** | **0.698** | **0.837** | 8M |
| | | 0.188 | 0.309 | 0.668 | 0.821 | 35M |
| | | 0.198 | 0.308 | 0.650 | 0.811 | 650M |
